# Cryptic, solo acylhomoserine lactone synthase from predatory myxobacterium suggests beneficial contribution to prey quorum signaling

**DOI:** 10.1101/849075

**Authors:** Hanan Albataineh, Maya Duke, Sandeep K. Misra, Joshua S. Sharp, D. Cole Stevens

## Abstract

Considered a key taxon in microbial communities, myxobacteria exist as coordinated swarms that utilize an excreted combination of lytic enzymes and specialized metabolites to facilitate predation of numerous microbial phyla. This capacity to produce biologically active metabolites and the associated abundance of natural product biosynthetic pathways contained within their genomes have motivated continued drug discovery efforts from myxobacteria. Of all the biosynthetic gene clusters associated with myxobacteria deposited in the antiSMASH database (∼1,000 total), only one putative acylhomoserine lactone synthase, *agpI*, was observed in genome data from the myxobacterium *Archangium gephyra*. Without an acylhomoserine lactone (AHL) receptor also apparent in the genome of *A. gephyra*, we sought to determine if AgpI was the first example of an orphaned AHL synthase. Herein we report the bioinformatic assessment of AgpI and discovery of a second myxobacterial AHL synthase from *Vitiosangium* sp. strain GDMCC 1.1324. During axenic cultivation conditions, no detectible AHL metabolites were observed in *A. gephyra* extracts. However, heterologous expression of each synthase in *Escherichia coli* provided detectible quantities of 3 AHL signals including 2 known AHLs, C8-AHL and C9-AHL. These results suggest that *A. gephyra* AHL production is dormant during axenic cultivation conditions and requires an unknown external cue for activation. The orphaned AHL synthase, AgpI, is the first to be reported from a predatory myxobacterium, and predator production of prey quorum signals provides unique insight into interspecies crosstalk within polymicrobial communities.

**Importance:** The presence of orphaned quorum signal receptors and associated recognition and response to exogenous acylhomoserine lactone quorum signals observed in microbial communities provides evidence for small molecule-mediated interspecies interactions. While the high occurrence of orphaned AHL receptors from bacteria that do not produce cognate AHL signals suggests the involvement of AHL signals as a shared chemical resource in polymicrobial communities, no orphaned AHL synthases have been determined to be functional in a species without an associated AHL receptor. An orphan signal synthase from a predatory myxobacterium provides an alternative perspective on the evolution and benefits of quorum signaling systems within these communities.

---

Ubiquitous throughout soils and marine sediments, myxobacteria utilize cooperative features to facilitate uniquely social lifestyles and exhibit organized predation of microbial prey (1-3). Often attributed to their predatory capabilities, an extraordinary number of biologically active specialized metabolites have been discovered from myxobacteria (4-8). Interest in this chemical space and the therapeutic potential associated with each elucidated natural product has motivated significant efforts towards continued discovery. Our recent survey of the unexplored, biosynthetic gene clusters from myxobacteria included in the antiSMASH database determined that the potential for such discovery from cultivable myxobacteria remains high (9-12). An oddity reported by this survey was the presence of a solo acylhomoserine lactone (AHL) synthase within the genome of the myxobacterium *Archangium gephyra* (9, 13, 14). As obligate cooperators numerous signaling systems have been associated with the coordination of myxobacterial motility and predation including A-signal, a quorum-like signal (1, 3, 15-17). However, no myxobacteria have been observed to produce AHL quorum signals.

Acylhomoserine lactone quorum signaling (QS) systems are abundant throughout Proteobacteria at-large (18). Considered autoinducers, AHLs bind to LuxR-type receptors which in turn activate LuxI-type AHL synthases (18). While a recent assessment of LuxR receptors included within or nearby specialized metabolite biosynthetic gene clusters (BGCs) reported the presence of a putative LuxR receptor from the marine myxobacterium *Haliangium ochraceum* DSM 14365, no AHL quorum signals or functional LuxI-type AHL synthases have been reported from myxobacteria (19). Intriguingly, the model myxobacterium *Myxococcus xanthus* demonstrates enhanced predatory features when exposed to a variety of exogenous AHLs despite having no obvious LuxR receptor within its genome (20). This phenomenon, often referred to as “eavesdropping,” has become a generally accepted cornerstone in hypotheses surrounding interspecies cross talk within polymicrobial communities, and the presence of solo or orphan LuxR receptors from species that do not produce AHL signals supports such communication (20-27). Putative solo-LuxR transcription factors with no accompanying LuxI synthases account for the majority of annotated LuxR proteins (18). However, as with *M. xanthus* (20), there are no LuxR receptors apparent in the genome of *A. gephyra*. This suggests that the observed LuxI-type synthase, AgpI, from *A. gephyra* is a solo-LuxI synthase. Considering the abundance of AHL QS systems throughout Proteobacteria other than myxobacteria, the uniqueness of this AHL synthase from *A. gephyra*, and the generalist diet of predatory myxobacteria that includes large swaths of AHL signaling proteobacteria, we hypothesize *agpI* was acquired horizontally (3, 28-30). Conversely, the benefit AHL production might provide a predatory myxobacterium remains non-obvious. Herein we report bioinformatic analysis, functional assessment, and heterologous expression of the myxobacterial AHL synthase AgpI.

## Results

### AgpI is highly homologous to functional AHL synthases

Located in the 20.6kb BGC referenced as cluster 33 from *A. gephyra* (NZ_CP011509) in the antiSMASH database (version 4.2.1) the 210aa gene product, AgpI (WP_047862734.1), is annotated as a putative autoinducer synthesis protein homologous to the GNAT family *N*-acetyltransferase, LuxI class of AHL synthases (10, 11, 21). None of the other annotated features neighboring *agpI* are obviously associated with AHL quorum signaling systems (Figure 1). Assessment of highly homologous LuxI synthases provided a second putative AHL synthase within the genome of the myxobacterium *Vitiosangium* sp. GDMCC 1.1324, deemed VitI (WP_108069305.1), with 98% coverage and 68.12% identity when comparing amino acid sequence data with AgpI (31). The absence of genome data for *V*. sp. in version 4.2.1 of the antiSMASH database explains the omission of this putative AHL synthase from our previous survey of myxobacterial biosynthetic space. The next highest scoring sequence from this analysis a GNAT family *N*-acetyltransferase (WP_055459978.1) from *Chelatococcus sambhunathii* has 96% coverage and 56.44% identity with AgpI (32, 33). Alignment and phylogenetic analysis of AgpI and VitI against an assortment of 17 LuxI-type synthases experimentally validated to produce AHL QS molecules, suggests common ancestry with the AHL synthases LuxI, LasI, and TraI from *Aliivibrio fischeri, Pseudomonas aeruginosa*, and *Rhizobium radiobacter* thus supporting our hypothesis that AgpI was horizontally acquired (Figure 2A and 2B) (34-45). Utilizing the genomic enzymology web tool EFI-EST developed by the Enzyme Function Initiative (EFI) to construct a sequence similarity network (SSN) that included 1,001 homologous entities as nodes and 124,346 edges, both AgpI and VitI are included in the central cluster family that contains the vast majority of homologous LuxI-type AHL synthases (Figure 2C) (46). From these data we conclude that both AgpI and VitI are likely AHL synthases as originally predicted by antiSMASH analysis. We also suggest that the shared ancestry observed from phylogenetic analysis and general absence of LuxI synthases from other myxobacterial phyla supports our hypothesis that these synthases were horizontally acquired.

**Figure 1:**
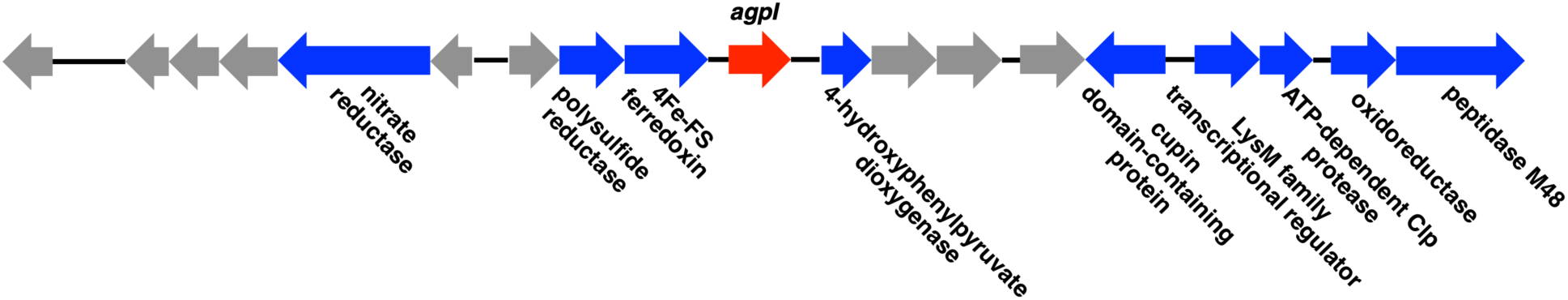
Cluster 33 from *A. gephyra* deposited in the antiSMASH database which includes the putative AHL synthase, *agpI*. All annotations included in the antiSMASH database provided and all hypothetical features are in grey (10, 11).

**Figure 2:**
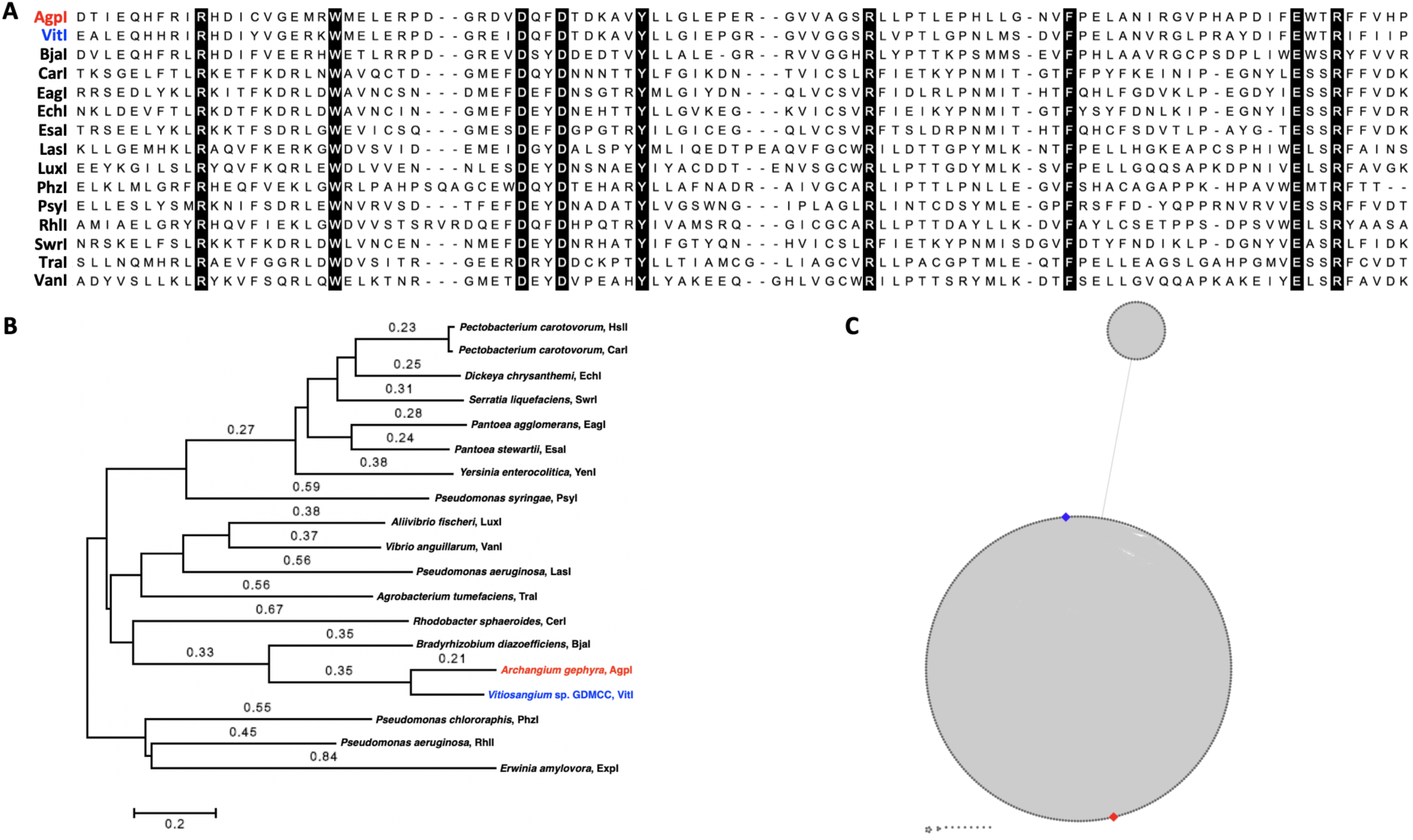
A) Alignment of LuxI synthases including AgpI and VitI with conserved residues boxed in black and B) a Minimum Evolution tree including AgpI and VitI rendered in MEGA7 using ClustalW aligned with AHL synthases experimentally confirmed to produce AHLs (68). Branch lengths ≤ 0.2 not depicted. C) Sequence similarity network rendered by EFI-EST analysis of AgpI amino acid sequence data with AgpI (red diamond) and VitI (blue diamond) indicated (46). To reduce complexity all nodes with ≥90% sequence similarity are represented as an individual aggregate node.

### Absence of a cognate AHL receptor in the genome of *A. gephyra*

While no obvious AHL-binding LuxR homolog was identified in the chromosome of *A. gephyra*, we sought to determine the presence of any potential AHL-binding domain using the conserved sequence for autoinducer binding domains (PF03472). Utilizing the blastp suite at NCBI, we assessed all 3,014 domains within the Pfam database classified as autoinducer binding domains for homology against the deposited genome of *A. gephyra* (14, 47). No features within the proteome of *A. gephyra* were sufficiently homologous to be considered to include an autoinducer binding domain. We next queried the Hidden Markov Model (HMM) associated with autoinducer binding domains deposited in Pfam against the proteome of *A. gephyra* using HMMSEARCH (supplemental data)(48, 49). The most significant hit (E-value 0.0015) a PAS domain S-box-containing protein also annotated as a GAF-domain-containing protein (WP_053066299.1) does not include significant sequence homology with LuxR-type, AHL receptors. Interestingly, similar analysis of *V*. sp. GDMCC 1.1324 provided a highly homologous LuxR-type receptor (WP_108076247.1). While the AHL receptor identified in the genome of *V*. sp. is not clustered near *vitI* as is typical of LuxI-LuxR type synthase-receptor pairs, we cannot assume both are unpaired orphans and instead consider VitI might not be a truly solo AHL synthase. From these data we determined AgpI to be an orphaned AHL synthase without any cognate LuxR receptor present in the genome of *A. gephyra*.

### *A. gephyra* does not produce AHLs during axenic cultivation

Cultivation of *A. gephyra* on VY/2 agar plates at 30°C for 21 days provided fully developed, wispy myxobacterial swarms encompassing the entirety of the plate surface. Homogenized agar and cellular contents were extracted using traditional organic phase techniques to provide extracts for LC-MS/MS analysis. The resulting datasets from LC-MS/MS analysis of *A. gephyra* extracts were analyzed against datasets generated from analytical standards for a variety of AHLs including C6-AHL, 3-oxo-C6-AHL, C8-AHL, and C11-AHL to determine the presence of any produced AHL-like metabolites. Data from resulting mass spectra were scrutinized using the Global Natural Products Social Molecular Networking (GNPS) platform to generate molecular networks depicting similarities in detected metabolite scaffolds inferred from ionized fragment commonalities (50). No metabolites that included the diagnostic AHL fragments at 102.0547 m/z and 74.0599 m/z associated with the core homoserine lactone moiety were detected in extracts from *A. gephyra* (51, 52). This data supports any one of the following conclusions *A. gephyra* does not produce AHL-like metabolites when grown axenically but may be active under other growth conditions; metabolites produced by AgpI do not possess structural similarity with typical AHL metabolites; or AgpI is simply nonfunctional. Cryptic or dormant BGCs are commonly observed during natural product discovery efforts, and various strategies to active cryptic BGCs have been developed including addition of exogenous chemical elicitors and heterologous expression of cryptic BGCs in an alternative host (53-55).

### Exogenous AHLs do not activate AgpI

Considering the typical autoinduction of LuxI synthases (18), we sought to determine if exogenous AHL metabolites might serve as elicitors and induce AgpI to provide observable AHL-like metabolites from *A. gephyra*. Experiments introducing the deuterated AHLs *N*-hexanoyl-L-homoserine lactone-*d*3 (C6-AHL-*d*3) and *N*-butyryl-L-homoserine lactone-*d*5 (C4-AHL-*d*5) to *A. gephyra* plates at 30 μM after 2 weeks of growth at 30°C were conducted to determine if exogenous AHLs induce AgpI activity. Deuterium-labelled analogs of C6-AHL and C4-AHL were utilized to provide the ability to decouple exogenous signals from structurally similar AHLs potentially induced by exogenous AHL introduction. However, using LC-MS/MS and molecular networking as previously described to analyze resulting extracts after an additional 7 days of growth following deuterated AHLs no metabolites possessing the core homoserine lactone moiety were detected in the deuterated AHL-exposed extracts from *A. gephyra* suggesting that AgpI activity is not induced by exogenous AHLs. While only 2 AHL signals were investigated, we suspect the likelihood of AHL autoinduction of AgpI is also diminished by the absence of a LuxR transcription factor typically responsible for activation of LuxI synthases.

### Heterologous expression of AgpI confirms functional production of AHLs

To explore the functionality of both AgpI and VitI and assumed biosynthesis of AHL-like metabolites, inducible codon-optimized constructs of *agpI* and *vitI* included in replicating plasmids suitable for expression in *Escherichia coli* were purchased. Heterologous expression of AgpI and VitI, subsequent extraction, LC-MS/MS analysis, and evaluation of molecular networks rendered by GNPS as previously described, provided a cluster family including 2 of 3 total nodes identified as C8-AHL (228.159 m/z) and C9-AHL (242.174 m/z) from internal GNPS public datasets as well as a third AHL metabolite detected at 226.144 m/z (Figure 3) (50). This cluster family was identical in both heterologous expression experiments suggesting that AgpI and VitI produce the same 3 AHL metabolites when heterologously expressed in *E. coli* with similar detected intensities for each AHL. Both C8-AHL and C9-AHL were confirmed to be present in AgpI and VitI extracts using analytical standards. Based on associated intensities, C8-AHL was the most abundant and the metabolite detected at 226.144 m/z was the least abundant AHL. No AHL-like entities were detected in control extracts from *E. coli* containing an empty pET28b expression plasmid. From the mass difference between C8-AHL and the unknown AHL detected at 226.144 m/z (2.015 Da measured vs. 2.01565 theoretical), as well as shared fragmentation patterns, we suggest the metabolite detected at 226.144 m/z to likely be an unsaturated analog of C8-AHL (Figure 4). From these experiments we determined that both AgpI and VitI are functional AHL synthases capable of producing the previously characterized AHLs C8-AHL and C9-AHL. These results also suggest *A. gephyra* produces AHLs and likely requires environmental cues or specific nutrients not present during our axenic cultivation conditions.

**Figure 3:**
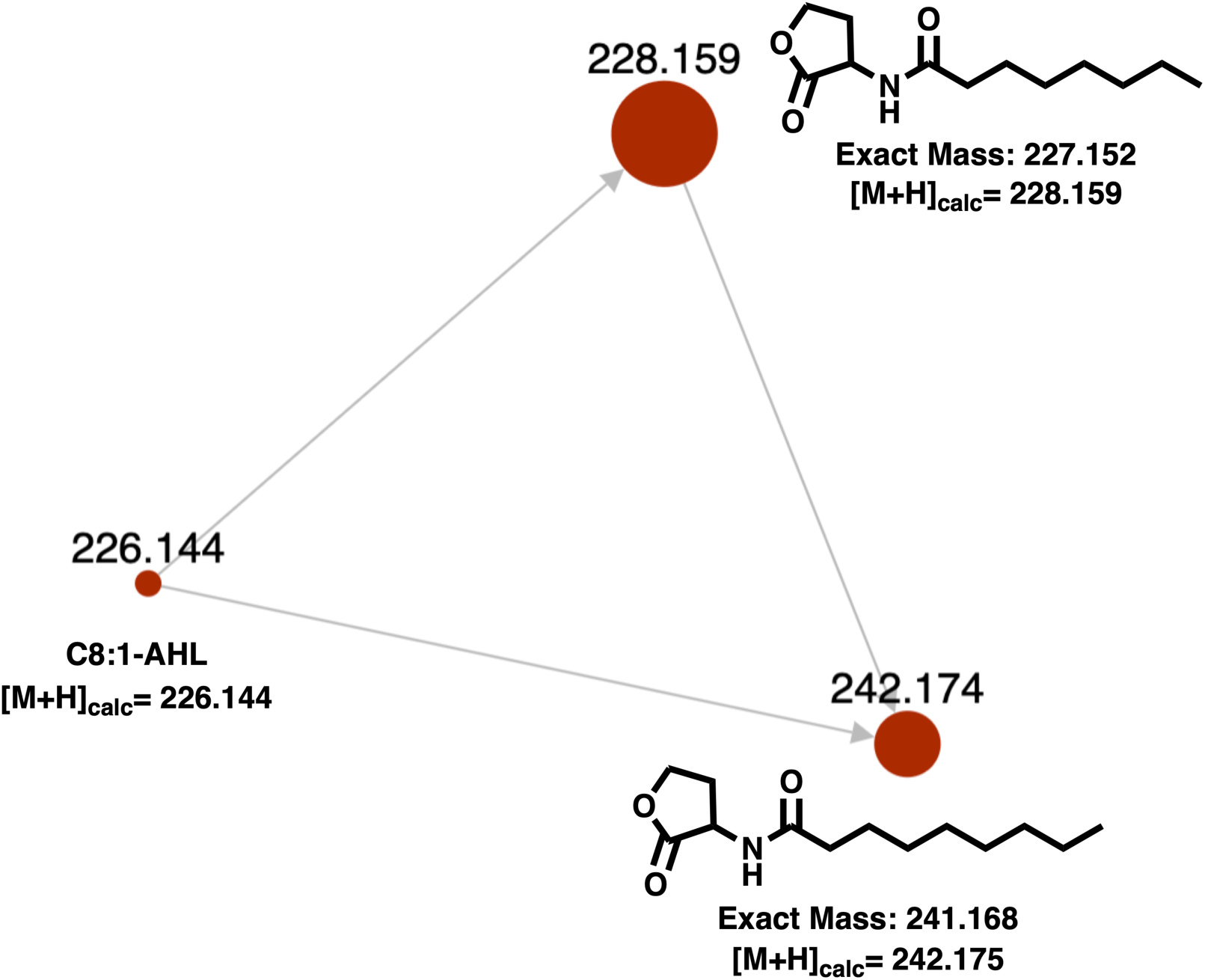
Molecular family from the molecular network of LC-MS/MS datasets from extracts of heterologous *E. coli* expressing AgpI rendered by GNPS (50). Detected m/z values from raw data positioned over each node with node diameter depicting associated intensities for each AHL.

**Figure 4:**
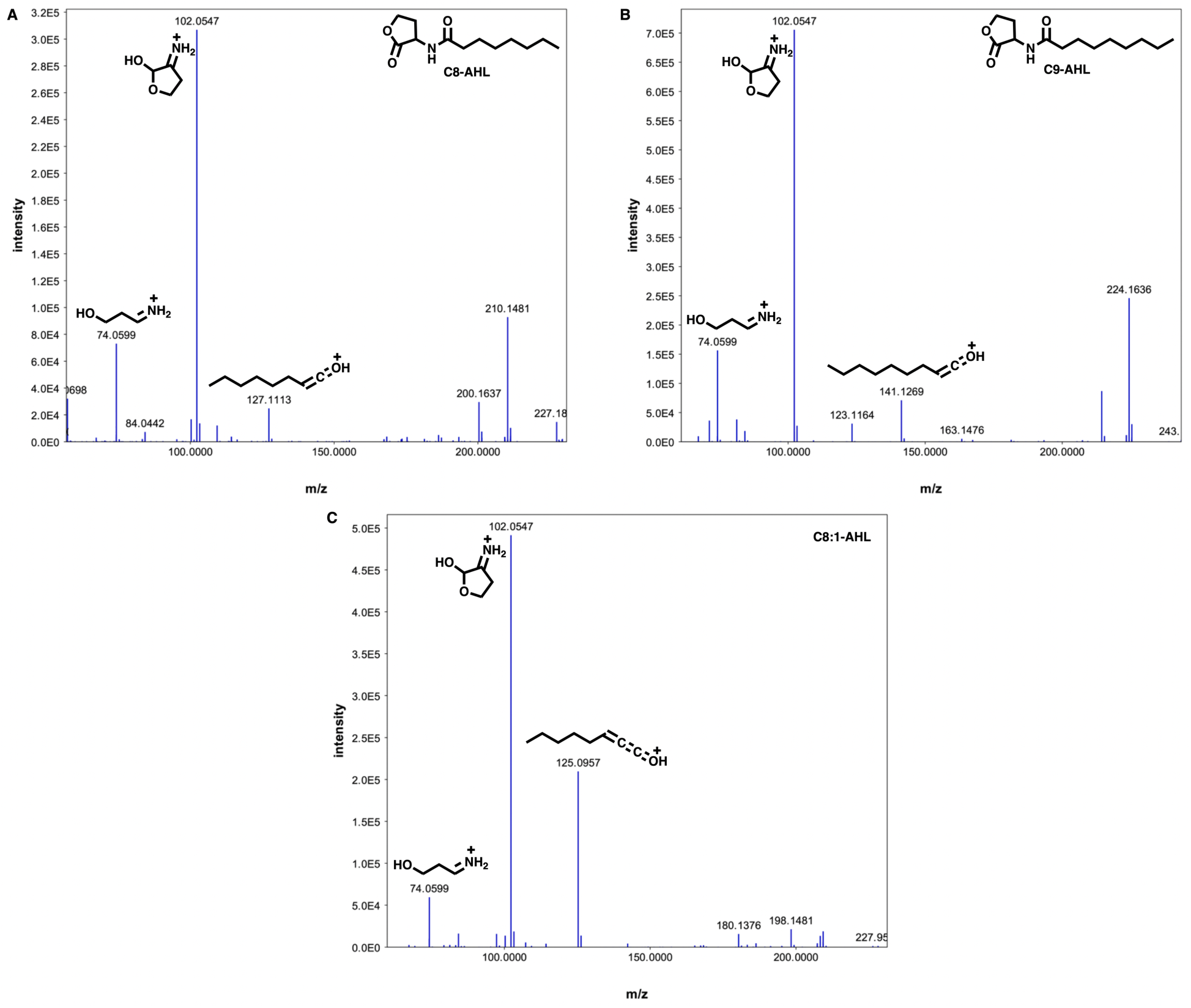
MS/MS fragmentation spectra with diagnostic fragments indicated for each AHL detected in extracts from heterologous *E. coli* expressing AgpI (51, 52).

## Discussion

Ultimately we conclude that the myxobacteria *A. gephyra* and *V*. sp. possess functional AHL synthases that produce the AHL signals C8-AHL and C9-AHL when heterologously expressed in *E. coli*. Considering the strong precedent for heterologous expression of AHL synthases in *E. coli* to determine produced AHL metabolites, we suggest that both *A. gephyra* and *V*. sp. capably produce one or all of the observed AHL signals and that AgpI is merely silent or cryptic during axenic cultivation of *A. gephyra* (56-61). However, we should also consider that these synthases could instead utilize an acyl-ACP precursor not available to the heterologous *E. coli* host, and we are actively exploring cultivation conditions that might induce native AHL production from *A. gephyra* (60, 61). While numerous bacteria have been observed to possess orphaned LuxR-type AHL receptors, production of AHL metabolites from a solo LuxI synthase without any cognate LuxR receptor also present in the genome is the first to be reported (22, 24, 26, 62). Although a functional orphaned LuxI-type synthase capable of producing AHLs has been reported from the sponge symbiont *Ruegeria* sp. KLH11, the strain also harbors 2 pairs of clustered LuxI/LuxR homologues (26, 63). We suggest that production of AHL quorum signals by myxobacteria supports the theoretical benefits of interspecies cross talk similar to functional, solo LuxR receptors (24, 64-66). We also propose that the more typical abundance of orphan LuxR receptors compared to the seemingly exceptional solo LuxI synthase reported here might correlate with the rarity of cooperative generalist predators (22, 30). The absence of any AHL metabolites in *A. gephyra* extracts suggests AgpI is cryptic or silent during axenic cultivation conditions and an unknown regulatory mechanism independent from a LuxR receptor to be involved. However, previously reported eavesdropping by *M. xanthus* and response to exogenous AHLs despite the absence of any AHL receptor with homology to LuxR suggests myxobacteria may possess an undiscovered, alternative means of AHL detection and response (20). While the benefit afforded predatory myxobacteria remains unclear, production of AHL signals known to regulate QS-associated physiological functions such as biofilm formation, specialized metabolism, and motility offers some insight (18). Predatory disruption of any one of these functions would likely benefit the fitness of *A. gephyra* by improving predation of quorum signaling prey. We consider these observations provide a unique perspective and support the continued investigation of small molecule interactions that contribute to microbial community structures and trophic levels.

## Materials and Methods

### Cultivation of *A. gephyra*

*Archangium gephyra* (DSM 2261) initially obtained from German Collection of Microorganisms in Braunschweig was grown on VY/2 agar (5 g/L baker’s yeast, 1.36 g/L CaCl_2_, 0.5 mg/L vitamin B_12_, 15 g/L agar, pH 7.2).

### Bioinformatic assessment of AgpI

The amino acid sequence for AgpI (WP_047862734.1) was submitted for blastp analysis and EFI-EST analysis (https://efi.igb.illinois.edu/efi-est/) using the default settings. Results from EFI-EST analysis were visualized using Cytoscape and are provided as supplemental data. Alignments from ClustalW and minimum evolution phylogenetic trees were rendered using MEGA7 (67, 68).

### Autoinducer binding site search

All 3,014 domains annotated as autoinducer binding domains (PF03472) deposited in Pfam were subjected to blastp analysis against the *A. gephyra* genome (NZ_CP011509.1). For HMMSEARCH analysis, the raw HHM for autoinducer binding domains was downloaded from Pfam (PF03472) and utilized as input for profile-HMM vs protein sequence database via HMMSEARCH with the taxonomy restrictions set to limit analysis to *A. gephyra* or *V*. sp. Results from this analysis are provided as supplemental data.

### Heterologous expression of AgpI and VitI in *E. coli*

Constructs of AgpI and VitI codon optimized for expression in *E. coli* situated in pET28b were purchased from Genscript (Piscataway, NJ). Sequence data for these constructs are provided as supplemental data. Heterologous host *E. coli* K207-3 was grown at 37**°**C in LB broth supplemented with 50μg/mL kanamycin, induced with 1μM IPTG at OD_600_=0.6, and grown overnight at 14**°**C to facilitate heterologous protein expression.

### Metabolite extraction and analysis

After 21 days of cultivation, *A. gephyra* plates were manually diced and extracted with excess EtOAc. Pooled EtOAc was filtered and dried *in vacuo* to provide crude extracts for LC-MS/MS analysis. Extracts from heterologous strains of *E. coli* were generated by Amberlite XAD-16 absorber resin facilitated extraction of clarified culture broths following cell lysis. LC-MS/MS analysis of the extracted samples was performed on an Orbitrap Fusion instrument (Thermo Scientific, San Jose, CA) controlled with Xcalibur version 2.0.7 and coupled to a Dionex Ultimate 3000 nanoUHPLC system. Samples were loaded onto a PepMap 100 C18 column (0.3 mm × 150 mm, 2 μm, Thermo Fisher Scientific). Separation of the samples was performed using mobile phase A (0.1% formic acid in water) and mobile phase B (0.1% formic acid in acetonitrile) at a rate of 6 μL/min. The samples were eluted with a gradient consisting of 5 to 60% solvent B over 15 min, ramped to 95 % B over 2 min, held for 3 min, and then returned to 5% B over 3 min and held for 8 min. All data were acquired in positive ion mode. Collision-induced dissociation (CID) was used to fragment molecules, with an isolation width of 3 m/z units. The spray voltage was set to 3600 volts, and the temperature of the heated capillary was set to 300 °C. In CID mode, full MS scans were acquired from m/z 150 to 1200 followed by eight subsequent MS2 scans on the top eight most abundant peaks. The orbitrap resolution for both the MS1 and MS2 scans was 120000. The expected mass accuracy was <3 ppm.

### Exogenous AHL exposure experiments

Stock solutions (10 mM) of *N*-hexanoyl-L-homoserine lactone-*d*3 (C6-AHL-*d*3) and *N*-butyryl-L-homoserine lactone-*d*5 (C4-AHL-*d*5) (Cayman Chemical) were prepared in DMSO. The required volumes of these stock solutions were filter sterilized and added to 14 days growing *A. gephyra* plates to give a final concentration of 30 μM. After 7 days of exogenous AHL exposure, *A. gephyra* plates were manually diced, extracted with excess EtOAc, and submitted to LC-MS/MS analysis as described before.

### GNPS dataset

Generated data were converted to .mzXML files using MS-Convert and mass spectrometry molecular networks were generated using the GNPS platform (http://gnps.ucsd.edu) (50). The corresponding Cytoscape file is provided as supplemental information. LC-MS/MS data for this analysis were also deposited in the MassIVE Public GNPS data set (MSV000084574).

## Supporting information

HMMsearch_Agephyra

HMMsearch_Vsp

blastP_Agephyra

AgpI_EFIEST_network_file

## Acknowledgements

The authors appreciate funding and support from the National Institute of Allergy and Infectious Diseases (R15AI137996), Fulbright (H.A), and the University of Mississippi.

## References

1. Cao P, Dey A, Vassallo CN, Wall D. 2015. How Myxobacteria Cooperate. J Mol Biol 427:3709–21.

2. Mohr KI. 2018. Diversity of Myxobacteria-We Only See the Tip of the Iceberg. Microorganisms 6.

3. Munoz-Dorado J, Marcos-Torres FJ, Garcia-Bravo E, Moraleda-Munoz A, Perez J. 2016. Myxobacteria: Moving, Killing, Feeding, and Surviving Together. Front Microbiol 7:781.

4. Bader CD, Panter F, Muller R. 2019. In depth natural product discovery - Myxobacterial strains that provided multiple secondary metabolites. Biotechnol Adv doi: 10.1016/j.biotechadv.2019.107480:107480.

5. Baltz RH. 2017. Gifted microbes for genome mining and natural product discovery. J Ind Microbiol Biotechnol 44:573–588.

6. Herrmann J, Fayad AA, Muller R. 2017. Natural products from myxobacteria: novel metabolites and bioactivities. Nat Prod Rep 34:135–160.

7. Korp J, Vela Gurovic MS, Nett M. 2016. Antibiotics from predatory bacteria. Beilstein J Org Chem 12:594–607.

8. Landwehr W, Wolf C, Wink J. 2016. Actinobacteria and Myxobacteria-Two of the Most Important Bacterial Resources for Novel Antibiotics. Curr Top Microbiol Immunol 398:273–302.

9. Gregory K, Salvador LA, Akbar S, Adaikpoh BI, Stevens DC. 2019. Survey of Biosynthetic Gene Clusters from Sequenced Myxobacteria Reveals Unexplored Biosynthetic Potential. Microorganisms 7.

10. Blin K, Medema MH, Kottmann R, Lee SY, Weber T. 2017. The antiSMASH database, a comprehensive database of microbial secondary metabolite biosynthetic gene clusters. Nucleic Acids Res 45:D555–D559.

11. Blin K, Pascal Andreu V, de Los Santos ELC, Del Carratore F, Lee SY, Medema MH, Weber T. 2019. The antiSMASH database version 2: a comprehensive resource on secondary metabolite biosynthetic gene clusters. Nucleic Acids Res 47:D625–D630.

12. Blin K, Shaw S, Steinke K, Villebro R, Ziemert N, Lee SY, Medema MH, Weber T. 2019. antiSMASH 5.0: updates to the secondary metabolite genome mining pipeline. Nucleic Acids Res 47:W81–W87.

13. Lang E, Schumann P, Tindall BJ, Mohr KI, Sproer C. 2015. Reclassification of Angiococcus disciformis, Cystobacter minus and Cystobacter violaceus as Archangium disciforme comb. nov., Archangium minus comb. nov. and Archangium violaceum comb. nov., unification of the families Archangiaceae and Cystobacteraceae, and emended descriptions of the families Myxococcaceae and Archangiaceae. Int J Syst Evol Microbiol 65:4032–4042.

14. Sharma G, Subramanian S. 2017. Unravelling the Complete Genome of Archangium gephyra DSM 2261T and Evolutionary Insights into Myxobacterial Chitinases. Genome Biol Evol 9:1304–1311.

15. Thiery S, Kaimer C. 2020. The Predation Strategy of Myxococcus xanthus. Front Microbiol 11:2.

16. Harris BZ, Kaiser D, Singer M. 1998. The guanosine nucleotide (p)ppGpp initiates development and A-factor production in myxococcus xanthus. Genes Dev 12:1022–35.

17. Kaplan HB, Kuspa A, Kaiser D. 1991. Suppressors that permit A-signal-independent developmental gene expression in Myxococcus xanthus. J Bacteriol 173:1460–70.

18. Papenfort K, Bassler BL. 2016. Quorum sensing signal-response systems in Gram-negative bacteria. Nat Rev Microbiol 14:576–88.

19. Brotherton CA, Medema MH, Greenberg EP. 2018. luxR Homolog-Linked Biosynthetic Gene Clusters in Proteobacteria. mSystems 3.

20. Lloyd DG, Whitworth DE. 2017. The Myxobacterium Myxococcus xanthus Can Sense and Respond to the Quorum Signals Secreted by Potential Prey Organisms. Front Microbiol 8:439.

21. Schuster M, Sexton DJ, Diggle SP, Greenberg EP. 2013. Acyl-homoserine lactone quorum sensing: from evolution to application. Annu Rev Microbiol 67:43–63.

22. Subramoni S, Venturi V. 2009. LuxR-family ‘solos’: bachelor sensors/regulators of signalling molecules. Microbiology 155:1377–85.

23. Subramoni S, Venturi V. 2009. PpoR is a conserved unpaired LuxR solo of Pseudomonas putida which binds N-acyl homoserine lactones. BMC Microbiol 9:125.

24. Wellington S, Greenberg EP. 2019. Quorum Sensing Signal Selectivity and the Potential for Interspecies Cross Talk. MBio 10.

25. Whiteley M, Diggle SP, Greenberg EP. 2017. Progress in and promise of bacterial quorum sensing research. Nature 551:313–320.

26. Tobias NJ, Brehm J, Kresovic D, Brameyer S, Bode HB, Heermann R. 2019. New vocabulary for bacterial communication. Chembiochem doi: 10.1002/cbic.201900580.

27. Hudaiberdiev S, Choudhary KS, Vera Alvarez R, Gelencser Z, Ligeti B, Lamba D, Pongor S. 2015. Census of solo LuxR genes in prokaryotic genomes. Front Cell Infect Microbiol 5:20.

28. Mendes-Soares H, Velicer GJ. 2013. Decomposing predation: testing for parameters that correlate with predatory performance by a social bacterium. Microb Ecol 65:415–23.

29. Morgan AD, MacLean RC, Hillesland KL, Velicer GJ. 2010. Comparative analysis of myxococcus predation on soil bacteria. Appl Environ Microbiol 76:6920–7.

30. Perez J, Moraleda-Munoz A, Marcos-Torres FJ, Munoz-Dorado J. 2016. Bacterial predation: 75 years and counting! Environ Microbiol 18:766–79.

31. Awal RP, Garcia R, Gemperlein K, Wink J, Kunwar B, Parajuli N, Muller R. 2017. Vitiosangium cumulatum gen. nov., sp. nov. and Vitiosangium subalbum sp. nov., soil myxobacteria, and emended descriptions of the genera Archangium and Angiococcus, and of the family Cystobacteraceae. Int J Syst Evol Microbiol 67:1422–1430.

32. Badhai J, Whitman WB, Das SK. 2016. Draft Genome Sequence of Chelatococcus sambhunathii Strain HT4T (DSM 18167T) Isolated from a Hot Spring in India. Genome Announc 4.

33. Panday D, Das SK. 2010. Chelatococcus sambhunathii sp. nov., a moderately thermophilic alphaproteobacterium isolated from hot spring sediment. Int J Syst Evol Microbiol 60:861–5.

34. White CE, Winans SC. 2007. Cell-cell communication in the plant pathogen Agrobacterium tumefaciens. Philos Trans R Soc Lond B Biol Sci 362:1135–48.

35. Schaefer AL, Greenberg EP, Oliver CM, Oda Y, Huang JJ, Bittan-Banin G, Peres CM, Schmidt S, Juhaszova K, Sufrin JR, Harwood CS. 2008. A new class of homoserine lactone quorum-sensing signals. Nature 454:595–9.

36. Rasch M, Andersen JB, Nielsen KF, Flodgaard LR, Christensen H, Givskov M, Gram L. 2005. Involvement of bacterial quorum-sensing signals in spoilage of bean sprouts. Appl Environ Microbiol 71:3321–30.

37. Pearson JP, Gray KM, Passador L, Tucker KD, Eberhard A, Iglewski BH, Greenberg EP. 1994. Structure of the autoinducer required for expression of Pseudomonas aeruginosa virulence genes. Proc Natl Acad Sci U S A 91:197–201.

38. Parsek MR, Schaefer AL, Greenberg EP. 1997. Analysis of random and site-directed mutations in rhII, a Pseudomonas aeruginosa gene encoding an acylhomoserine lactone synthase. Mol Microbiol 26:301–10.

39. Nhu Lam M, Dudekula D, Durham B, Collingwood N, Brown EC, Nagarajan R. 2018. Insights into beta-ketoacyl-chain recognition for beta-ketoacyl-ACP utilizing AHL synthases. Chem Commun (Camb) 54:8838–8841.

40. Milton DL, Hardman A, Camara M, Chhabra SR, Bycroft BW, Stewart GS, Williams P. 1997. Quorum sensing in Vibrio anguillarum: characterization of the vanI/vanR locus and identification of the autoinducer N-(3-oxodecanoyl)-L-homoserine lactone. J Bacteriol 179:3004–12.

41. Engebrecht J, Silverman M. 1984. Identification of genes and gene products necessary for bacterial bioluminescence. Proc Natl Acad Sci U S A 81:4154–8.

42. Eberl L, Winson MK, Sternberg C, Stewart GS, Christiansen G, Chhabra SR, Bycroft B, Williams P, Molin S, Givskov M. 1996. Involvement of N-acyl-L-hormoserine lactone autoinducers in controlling the multicellular behaviour of Serratia liquefaciens. Mol Microbiol 20:127–36.

43. Dong SH, Frane ND, Christensen QH, Greenberg EP, Nagarajan R, Nair SK. 2017. Molecular basis for the substrate specificity of quorum signal synthases. Proc Natl Acad Sci U S A 114:9092–9097.

44. Chakrabarti S, Sowdhamini R. 2003. Functional sites and evolutionary connections of acylhomoserine lactone synthases. Protein Eng 16:271–8.

45. Brader G, Sjoblom S, Hyytiainen H, Sims-Huopaniemi K, Palva ET. 2005. Altering substrate chain length specificity of an acylhomoserine lactone synthase in bacterial communication. J Biol Chem 280:10403–9.

46. Zallot R, Oberg N, Gerlt JA. 2019. The EFI Web Resource for Genomic Enzymology Tools: Leveraging Protein, Genome, and Metagenome Databases to Discover Novel Enzymes and Metabolic Pathways. Biochemistry 58:4169–4182.

47. El-Gebali S, Mistry J, Bateman A, Eddy SR, Luciani A, Potter SC, Qureshi M, Richardson LJ, Salazar GA, Smart A, Sonnhammer ELL, Hirsh L, Paladin L, Piovesan D, Tosatto SCE, Finn RD. 2019. The Pfam protein families database in 2019. Nucleic Acids Res 47:D427–D432.

48. Finn RD, Clements J, Eddy SR. 2011. HMMER web server: interactive sequence similarity searching. Nucleic Acids Res 39:W29–37.

49. Wheeler TJ, Eddy SR. 2013. nhmmer: DNA homology search with profile HMMs. Bioinformatics 29:2487–9.

50. Wang M, Carver JJ, Phelan VV, Sanchez LM, Garg N, Peng Y, Nguyen DD, Watrous J, Kapono CA, Luzzatto-Knaan T, Porto C, Bouslimani A, Melnik AV, Meehan MJ, Liu WT, Crusemann M, Boudreau PD, Esquenazi E, Sandoval-Calderon M, Kersten RD, Pace LA, Quinn RA, Duncan KR, Hsu CC, Floros DJ, Gavilan RG, Kleigrewe K, Northen T, Dutton RJ, Parrot D, Carlson EE, Aigle B, Michelsen CF, Jelsbak L, Sohlenkamp C, Pevzner P, Edlund A, McLean J, Piel J, Murphy BT, Gerwick L, Liaw CC, Yang YL, Humpf HU, Maansson M, Keyzers RA, Sims AC, Johnson AR, Sidebottom AM, Sedio BE, et al. 2016. Sharing and community curation of mass spectrometry data with Global Natural Products Social Molecular Networking. Nat Biotechnol 34:828–837.

51. Patel NM, Moore JD, Blackwell HE, Amador-Noguez D. 2016. Identification of Unanticipated and Novel N-Acyl L-Homoserine Lactones (AHLs) Using a Sensitive Non-Targeted LC-MS/MS Method. PLoS One 11:e0163469.

52. Gould TA, Herman J, Krank J, Murphy RC, Churchill ME. 2006. Specificity of acyl-homoserine lactone synthases examined by mass spectrometry. J Bacteriol 188:773–83.

53. Nguyen CT, Dhakal D, Pham VTT, Nguyen HT, Sohng JK. 2020. Recent Advances in Strategies for Activation and Discovery/Characterization of Cryptic Biosynthetic Gene Clusters in Streptomyces. Microorganisms 8.

54. Ochi K, Hosaka T. 2013. New strategies for drug discovery: activation of silent or weakly expressed microbial gene clusters. Appl Microbiol Biotechnol 97:87–98.

55. Xu F, Nazari B, Moon K, Bushin LB, Seyedsayamdost MR. 2017. Discovery of a Cryptic Antifungal Compound from Streptomyces albus J1074 Using High-Throughput Elicitor Screens. J Am Chem Soc 139:9203–9212.

56. Zheng H, Zhong Z, Lai X, Chen WX, Li S, Zhu J. 2006. A LuxR/LuxI-type quorum-sensing system in a plant bacterium, Mesorhizobium tianshanense, controls symbiotic nodulation. J Bacteriol 188:1943–9.

57. Neumann A, Patzelt D, Wagner-Dobler I, Schulz S. 2013. Identification of new N-acylhomoserine lactone signalling compounds of Dinoroseobacter shibae DFL-12(T) by overexpression of luxI genes. Chembiochem 14:2355–61.

58. Dang HT, Komatsu S, Masuda H, Enomoto K. 2017. Characterization of LuxI and LuxR Protein Homologs of N-Acylhomoserine Lactone-Dependent Quorum Sensing System in Pseudoalteromonas sp. 520P1. Mar Biotechnol (NY) 19:1–10.

59. Britstein M, Devescovi G, Handley KM, Malik A, Haber M, Saurav K, Teta R, Costantino V, Burgsdorf I, Gilbert JA, Sher N, Venturi V, Steindler L. 2016. A New N-Acyl Homoserine Lactone Synthase in an Uncultured Symbiont of the Red Sea Sponge Theonella swinhoei. Appl Environ Microbiol 82:1274–1285.

60. Hoshino S, Onaka H, Abe I. 2019. Activation of silent biosynthetic pathways and discovery of novel secondary metabolites in actinomycetes by co-culture with mycolic acid-containing bacteria. J Ind Microbiol Biotechnol 46:363–374.

61. Mao D, Okada BK, Wu Y, Xu F, Seyedsayamdost MR. 2018. Recent advances in activating silent biosynthetic gene clusters in bacteria. Curr Opin Microbiol 45:156–163.

62. Chugani S, Greenberg EP. 2014. An evolving perspective on the Pseudomonas aeruginosa orphan quorum sensing regulator QscR. Front Cell Infect Microbiol 4:152.

63. Zan J, Choi O, Meharena H, Uhlson CL, Churchill ME, Hill RT, Fuqua C. 2015. A solo luxI-type gene directs acylhomoserine lactone synthesis and contributes to motility control in the marine sponge symbiont Ruegeria sp. KLH11. Microbiology 161:50–6.

64. Venturi V, Bertani I, Kerenyi A, Netotea S, Pongor S. 2010. Co-swarming and local collapse: quorum sensing conveys resilience to bacterial communities by localizing cheater mutants in Pseudomonas aeruginosa. PLoS One 5:e9998.

65. Noel JT, Joy J, Smith JN, Fatica M, Schneider KR, Ahmer BM, Teplitski M. 2010. Salmonella SdiA recognizes N-acyl homoserine lactone signals from Pectobacterium carotovorum in vitro, but not in a bacterial soft rot. Mol Plant Microbe Interact 23:273–82.

66. McDougald D, Srinivasan S, Rice SA, Kjelleberg S. 2003. Signal-mediated cross-talk regulates stress adaptation in Vibrio species. Microbiology 149:1923–33.

67. Sievers F, Higgins DG. 2018. Clustal Omega for making accurate alignments of many protein sequences. Protein Sci 27:135–145.

68. Kumar S, Stecher G, Tamura K. 2016. MEGA7: Molecular Evolutionary Genetics Analysis Version 7.0 for Bigger Datasets. Mol Biol Evol 33:1870–4.

